# Understanding the potential benefits of adaptive therapy for metastatic melanoma

**DOI:** 10.1101/2020.10.16.343269

**Authors:** Eunjung Kim, Joel S. Brown, Zeynep Eroglu, Alexander R.A. Anderson

## Abstract

Adaptive therapy is an evolution-based treatment approach that aims to maintain tumor volume by employing minimum effective drug doses or timed drug holidays. For successful adaptive therapy outcomes, it is critical to find the optimal timing of treatment switch points. Mathematical models are ideal tools to facilitate adaptive therapy dosing and switch time points. We developed two different mathematical models to examine interactions between drug-sensitive and resistant cells in a tumor. The first model assumes genetically fixed drug-sensitive and resistant populations that compete for limited resources. Resistant cell growth is inhibited by sensitive cells. The second model considers phenotypic switching between drug-sensitive and resistant cells. We calibrated each model to fit melanoma patient biomarker changes over time and predicted patient-specific adaptive therapy schedules. Overall, the models predict that adaptive therapy would have delayed time to progression by 6-25 months compared to continuous therapy with dose rates of 6%-74% relative to continuous therapy. We identified predictive factors driving the clinical time gained by adaptive therapy. The first model predicts 6-20 months gained from continuous therapy when the initial population of sensitive cells is large enough, and when the sensitive cells have a large competitive effect on resistant cells. The second model predicts 20-25 months gained from continuous therapy when the switching rate from resistant to sensitive cells is high and the growth rate of sensitive cells is low. This study highlights that there is a range of potential patient specific benefits of adaptive therapy, depending on the underlying mechanism of resistance, and identifies tumor specific parameters that modulate this benefit.

## Introduction

Current targeted therapies in melanoma are based on continuous treatment. Patients with advanced BRAFV600E mutant melanoma are eligible for targeted therapy with BRAF and MEK-inhibitors, with objective tumor response rates in up to 75% of the patients (1, 2). Despite impressive initial responses, a majority of patients with metastatic melanoma experience disease progression. Median progression-free survival ranges from 11-15 months (3, 4). In melanoma, a major driver of this resistance is intratumor heterogeneity (5). Adaptive therapy is an evolutionary inspired treatment strategy that exploits this heterogeneity, specifically harnessing competition between different cancer cell types for limited resources (6, 7). Standard treatments using maximum tolerated dose (MTD) rapidly remove drug-sensitive populations. If drug-resistant cells are present before treatment (8-10), then this aggressive MTD treatment alters the competition between drug-sensitive cells with drug-resistant populations. MTD treatments release resistant cells from competition with their sensitive counterparts via competitive release. As a result, the resistant cells rapidly come to dominate the tumor (11, 12). Several studies have shown that phenotypic switching between sensitive and resistance types provides another avenue for acquired resistance during therapy (9, 13, 14). Once resistant cells dominate, treatment fails since it no longer prevents tumor growth. The goal of adaptive therapy is to control each patient’s tumor volume by allowing a significant number of sensitive cells to survive through treatment breaks or dose-modulation. The central mechanism of adaptive therapy is competition, in that any remaining sensitive cells can compete with the resistant population, effectively suppressing their growth and significantly extending progression free survival (15-17).

Adaptive therapy has successfully controlled cancers in preclinical xenograft model systems (18, 19). An ongoing metastatic castrate-resistant prostate cancer clinical trial at Moffitt Cancer Center (NCT02415621) has shown that adaptive therapy can delay disease progression for 27 months using just 53% cumulative drug rate compared to standard of care (MTD) (20). For melanoma, previous work has shown that intermittent dosing of a BRAF inhibitor, vemurafenib, given to patient-derived mouse model systems, controlled tumor volumes significantly better than continuous therapy (21). But this fixed intermittent strategy is less likely to be effective in patients due to inter-patient heterogeneity. In fact, a trial that explored fixed 5 weeks on/3 weeks off dosing of a BRAF/MEK inhibitor regimen, dabrafenib + trametinib, demonstrated a lower median progression-free survival than standard, continuous dosing in patients with metastatic melanoma (22). Our group has previously shown that a mathematically driven adaptive approach can work *in vivo* using xenograft mouse models of melanoma (23). We posit that this personalized treatment approach, whereby mathematical models facilitate optimizing a patient’s treatment regime based on their current tumor state and historic response, will be far more effective.

Several challenges exist in designing adaptive therapies. First, optimizing the timing of treatment withdrawal and re-challenge for each patient is difficult to implement in clinical practice. Second, it is critical to identify predictive factors for selecting patients who will likely benefit the most from this adaptive therapy. The effectiveness of adaptive therapy will vary among patients, as observed in the prostate cancer trial (20). Several mathematical models have already been developed to address these challenges. Combined with *in vitro* experiments, an agent-based model predicted that spatially constrained competition drove an effective control of resistant cell growth (15). Another agent based model developed by Gallaher et al. assumed a cost of resistance, with resistant cells having a slower growth rate in the absence of treatment (16). They modeled resistance as a continuous phenotypic trait and revealed three key factors for successful adaptive therapy: the proportion of resistant cells prior at the start of treatment, the rate of cancer cell migration, and the speed of evolution towards more resistant phenotypes. An ordinary differential equation model developed by Hansen et al. identified thresholds for the level of initial resistance for successful adaptive therapy (17). West et al. developed an evolutionary game theory model to determine the optimal timing of multi-drug adaptive therapies (24). Brady-Nicholls et al. developed patient-specific mathematical models that explain prostate cancer inter-tumor dynamics that can guide intermittent androgen deprivation therapy (25). Mathematical modeling has shown the consequences of having spontaneous versus induced resistance responses to therapy (26). Strobl et al. showed how the turnover rate of cancer cells within the tumor influences benefits derived from adaptive therapy (27). More general and deeper mathematical analysis by Viossat and Noble predict the clinical benefits of tumor containment therapy (28).

Here we present two different ordinary differential equation models that assume different modes of competition between sensitive and resistant populations. We then calibrate both models to a cohort of 8 patients with metastatic melanoma. By simulating adaptive therapy schedules with these models, we identify predictive factors that correlate with the largest clinical gains compared to continuous MTD therapy. The first model is a Lotka-Volterra competition model (LV) (29), where genetically fixed drug-sensitive and -resistant populations compete for limited resources. The model assumes resistant cell growth inhibition by a drug-sensitive population. The second model considers phenotypic switching (SW). In addition to competition within and between cell phenotypes, this model allows for switching from sensitive to resistant states or vice-versa when treatment is on or off, respectively. We estimated model parameters by minimizing the difference between model prediction and patient data. For each patient, we used longitudinal data from a serologic marker that is used to monitor advanced melanoma.

The cohort of patients had advanced/metastatic melanoma. All were treated with continuous therapy at MTD. Their therapy consisted of BRAF/MEK inhibitors (either vemurafenib + cobimetinib, or dabrafenib + trametinib). Several had disease progression within 6 months of treatment (Figure 1). Whilst melanoma does not have an ideal biomarker of burden, LDH (Lactate dehydrogenase) is clinically used in melanoma treatment decision making as a correlate of tumor burden and cancer dynamics. LDH is the only serologic marker used for monitoring advanced melanoma in the US (30). Elevated serum LDH is associated with worse outcomes in patients treated with BRAF/MEK inhibitors, based on the results of a pooled analysis of three trials involving dabrafenib/trametinib with over 600 patients (31). In the cohort of the current study, all patients had an elevated LDH at the start of treatment, and serial LDH levels were measured in blood at baseline and during routine blood draws approximately every 2-4 weeks (Fig. S1 & Table S1). LDH was used as a biomarker to correlate with melanoma tumor burden, which could only be measured directly in patients with computed tomography (CT) imaging every 2 to 3 months. It is worth noting that an elevated level of LDH appeared to correlate with disease response and progression determined by CT imaging (Fig. S1 & Table S1). Temporal LDH profiles for each of the 8 patients highlight heterogeneous responses, with some rapidly developing therapy resistance (P6-8), while others took longer to progress (P1-5). Note that the initial tumor burden level is also different across the patients (LDH range: ~300 to ~1500).

**Figure 1.**
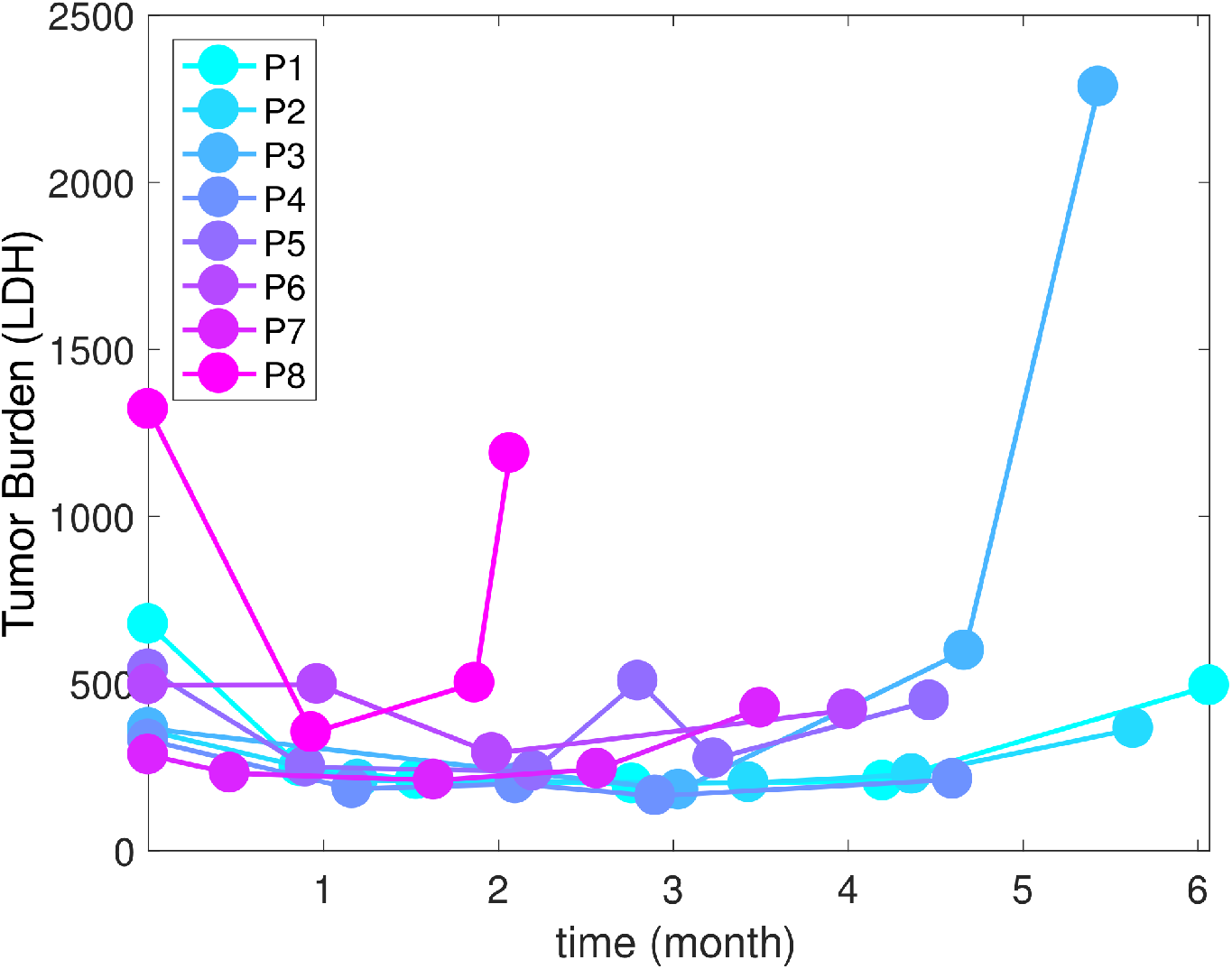
The tumor burdens (measured by LDH level units/liter) of 8 melanoma patients in months since the beginning of therapy. All 8 patients received continuous MTD of targeted therapy (BRAF & MEK inhibitor).

Model calibration (see methods) to the melanoma patient LDH data provided a suite of parameter sets that fit the patient data equally well, defining a virtual cohort of patients (32). Using this virtual cohort, we used the models to predict what might have been the patient’s responses to different adaptive therapy schedules. Our results show that adaptive therapies can delay the time to progression up to several months with less cumulative drug dose rate compared to the continuous MTD regime. Further, we identified key model parameters that determine the benefit of adaptive therapy that could be used to select patients suitable for this evolutionary therapeutic approach.

### Mathematical modeling

The first model uses Lotka-Volterra (LV) competition equations to describe the competition between two distinct cancer cell populations, drug-sensitive (*S*) and drug-resistant (*R*) cells (Fig. 2A):

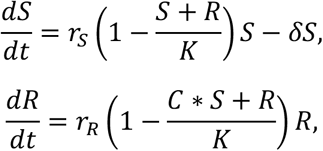

where *r_S,R_* indicate the intrinsic growth rates of *S* and *R*, respectively. The term *δ* > 0 imposes a death rate on *S* due to therapy. In the absence of treatment, we set *δ* = 0. Furthermore, we assume that treatment stops any proliferation by sensitive cells, and so we set *r_S_* = 0 when treatment is on and *r_S_* > 0 when treatment is off. The two populations, *S* and *R*, share the same carrying capacity *K*, the maximum size of the tumor due to nutrient and space constraints. The coefficient *C* scales the degree to which sensitive cells inhibit the growth rate of resistant cells. If *C* > 1 (or *C* < 1), then sensitive cells have a greater (or smaller) competitive effect on resistant cells than resistant cells have on themselves.

**Figure 2.**
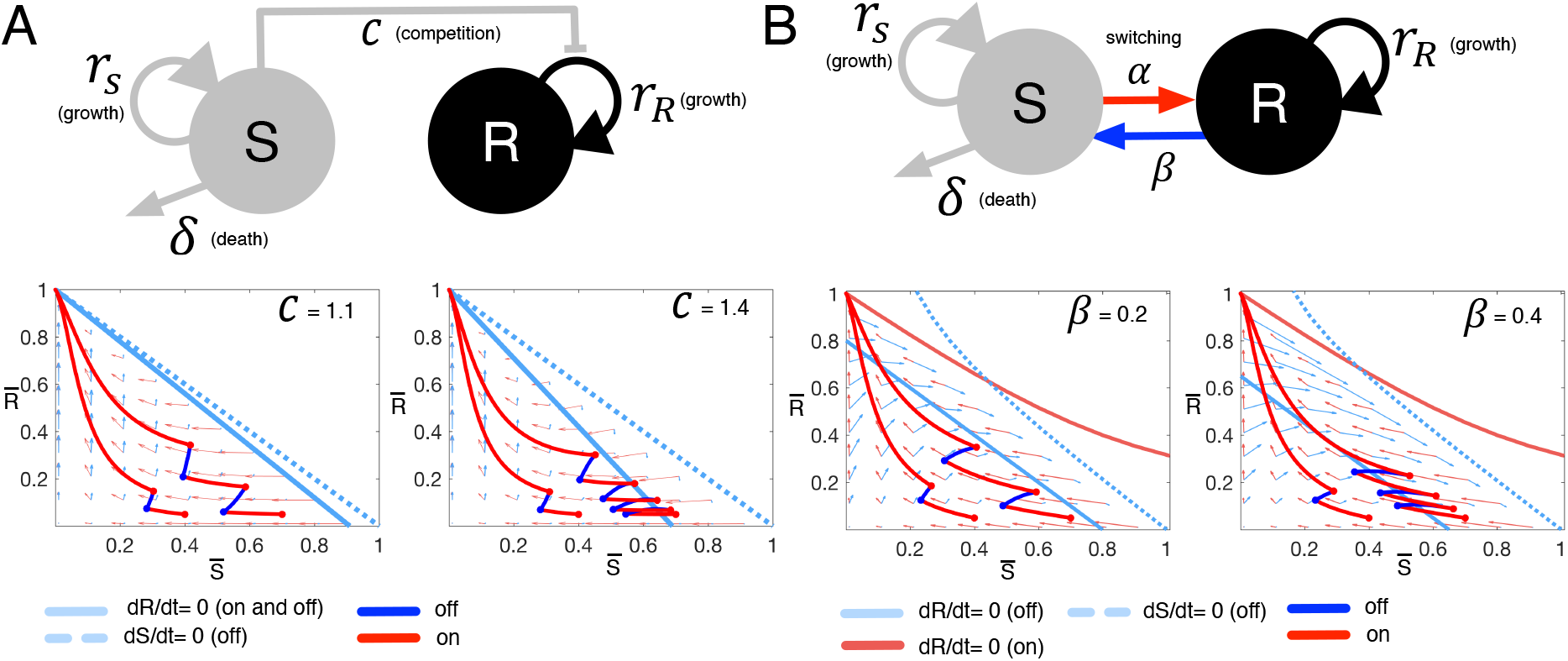
Tumor dynamics for the two mathematical models of tumor growth and treatment. A. Upper panel: LV model structure. S: drug-sensitive cancer cell population, outgoing arrow: death by a therapy, *R*: drug-resista nt cancer cell population, lin e to *R* from S: inhibition of *R* growth via competition, circular arrow: growth. Bottom pane I: temporal dynamics under i ntermittent therapy simulation with two different competition coefficients “*C* = 1.1 on the left & *C* = 1.4 on the right) and two different initial conditions (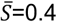 and 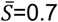 with 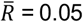 in both scenarios, 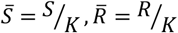). Coordinates: non-dimensionalized sensitive cell 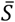 (horizontal axis) & resistant 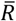 (vertical axis). (1,0) represents *S = K* & *R* = 0, (0,1) repre sents *S* = 0, & *R = K*. Rate of change of 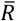 with respect to time is zero along the pale blue solid line when treatment is on and off. The rate of change of 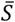 is zero along the pale blue dotted line when treatment is off. When treatment is on, the rate of change of 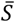 is zero along with y-axis. Temporal dynamics of the two cell types are traced along trajectories for intermittent treatment (red: on & blue: off). B. Upper panel: SW model structure. *S*: drug-sensitive cancer cell population, outgoing arrow: death by therapy, *R*: drug-resistant cancer cell population, a line from *S* to *R*: switching from *S* to *R* duri ng treatment on, a line from *R* to *S*: switching from *R* to *S* during treatment holidays, circular arrow: growth. Bottom panel: Temporal dynamics under intermittent th erapy simulation with two different *R*->*S* switching rates (*β*) and two d i fferent initial conditions (the same as for LV simulations). The rate of change of resistant cell population with respect to time is zero along two solid lines (pale blue: treatment off and pale red: treatment on). The rate of change of ! is zero along the pale blue dotted line when treatment is off.

As a second variant of the therapy model, we consider the case where phenotypic plasticity allows for switching between drug-sensitive and resistant cell types (SW, Fig. 2B):

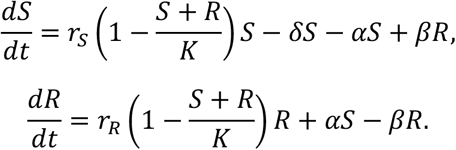

This model is a direct extension of the first model, and as such, all overlapping parameters are the same. There are two key differences. First, sensitive cells no longer have a scaled effect on resistant cells, and so *C* = 1. Second, sensitive cells can switch to resistant ones at rate *α*, or resistant cells to sensitive ones at rate *β*, depending on whether treatment is on or off. Note that *α* is non-zero when treatment is on and zero otherwise, whereas *β* is non-zero when treatment is off and zero when treatment is on.

To more easily visualize and understand the dynamics of these two models, we non-dimensionalized them 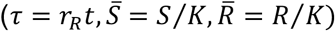 and examined their dynamics in the phase plane of 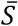 *vs*. 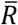. In the LV model, the competition coefficient *C* and initial populations 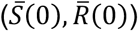 determine the number of intermittent therapy cycles before resistance dominates (Fig. 2A). Intermittent therapy results in more on-off treatment cycles when both *C* and 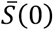 are large (Fig. 2A bottom panel). In the SW model (Fig. 2B), intermittent therapy results in more on-off treatment cycles when both the initial sensitive population 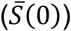 and the switching rate from resistant to sensitive populations (*β*) are large (Fig. 2B bottom panel). These analyses help focus our attention on the key parameters that might determine the efficacy of adaptive therapy.

### Parameter estimation

We identified model parameters that minimized the difference between model predictions and patient data (Figure 1). The cost function for this optimization is to minimize *L*_2_ norm of the difference,

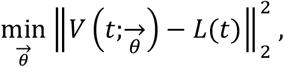

where 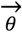 is the entire model parameter set, and *V* is the predicted total tumor burden (*V = S + R*) at time *t*, and *L* is actual tumor burden of each patient at time *t.* Here, we assume that LDH is directly proportional to *S + R*. The LV model parameter set includes intrinsic growth rates, carrying capacity, death rate, and the competition coefficient. Note, we assume that sensitive cells do not divide when treatment is on (*r_s_* = 0, with therapy on). All patient data is for continuous treatment (at the time of writing, we do not have intermittent therapy results for such patients). Thus, a parameter set for the LV model is *θ* = {*S*_0_, *K, δ, r_R_, C*}, where *S*_0_ is the initial number of sensitive cells and *R*_0_ = *LDH*_0_(1 − *S*_0_) is the initial number of resistant cells. In the SW model, the transition rate from resistant to sensitive is assumed to be zero when treatment is on. The parameter set for the SW model is *θ* = {*S*_0_, *K, δ, r_R_, α*}. We employed a steepest descent optimization algorithm with implicit filtering (33) to identify best-fit parameters for both models (Supplementary excel file). Parameter estimations were conducted in MATLAB using the implicit filtering algorithm (33). We generated patient-specific parameter estimates for each model. The fits to the individual patient data are shown in Fig. S2-3, Fig. 3A, & Fig. 3D. Across patients, under the LV model, time to progression is negatively correlated with the intrinsic growth rate of resistant cells (*r_R_*) and the competition coefficient (*C*) (Fig. 3B). Under the SW model, time to progression is negatively correlated with the intrinsic growth rate of resistant cells (Fig. 3E).

**Figure 3.**
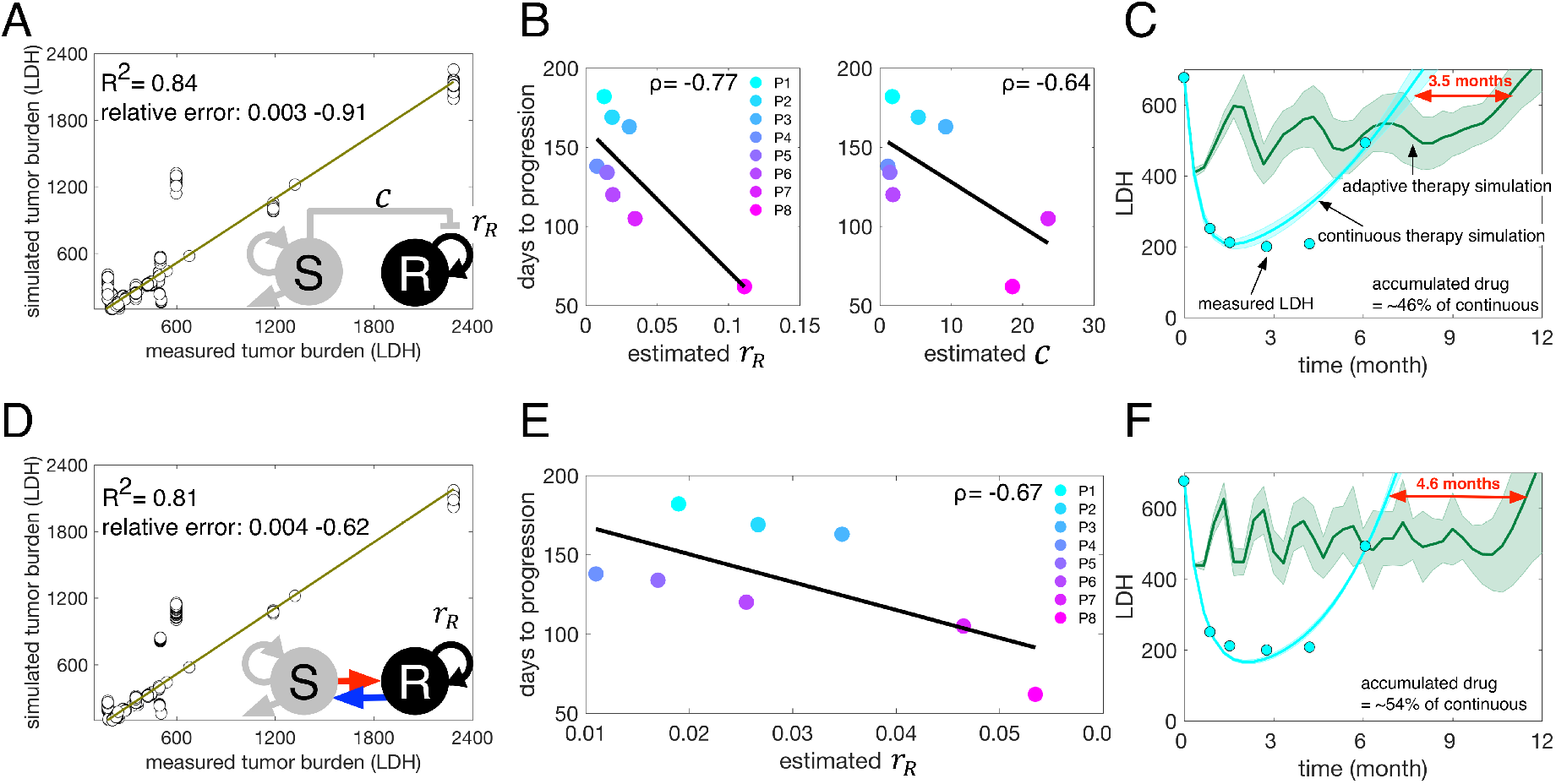
Mathematical model calibration & adaptive therapy simulation. A: LV model fits to the temporal LDH data from 8 patients. R^2^ = 0.84 when comparing the real patient LDH (horizontal axis) to the simulated tumor LDH (vertical axis). B: Time to progression under continuous therapy is negatively correlated with both the estimated intrinsic growth rate of resistant cells (*ρ* = −0.77) and the competition coefficient (*ρ* = −0.64). C: Adaptive therapy and continuous therapy simulation for patient #1 (P1) using the calibrated LV model. Cyan dots: LDH data of P1, cyan line: LV model fits. Green: predicted adaptive therapy (dark green: mean & shaded region: standard error). D: SW model fits to the temporal LDH data for the 8 patients. R^2^ = 0.81 when comparing the real patient LDH (horizontal axis) and simulated tumor LDH (vertical axis). E: Time to progression under continuous therapy is negatively correlated with the estimated intrinsic growth rate of resistant cells (*ρ* = −0.67). F: Adaptive therapy and continuous therapy simulation for patient 1 (P1) using the calibrated SW model. Cyan dots: P1 LDH data, cyan line: SW model fit. Green: predicted adaptive therapy (dark green: mean & shaded region: standard error).

## Results

### Adaptive therapy delays time to progression

Both the LV and SW models provide fits to the patient LDH data (Fig. 3A & D, *R*^2^ = 0.84 and *R*^2^ = 0.81, respectively, when comparing actual to predicted across all patient data points). Model parameterization generated multiple similar fits to the data with a similar error. Since there is no single ideal fit, we utilized the top 50 fits for each model and for each patient, resulting in a cohort of 400 virtual patients (8 patients, 50 virtual patients for each real patient) for each model, LV and SW, respectively). Using this virtual cohort, we simulated both continuous and adaptive therapy for each of the virtual patients. For adaptive therapy simulations, we assumed treatment dose per treatment day is the same as the dose in continuous therapy. We stopped treatment when a patient’s LDH level dropped to 50% of their initial LDH. Treatment remained off until the LDH grew back to its initial level. We estimated time to disease progression as the moment when a virtual patient’s LDH reached 150% of the patient’s initial LDH. For each virtual patient, we determined the time gained from adaptive therapy relative to continuous therapy by subtracting the time to progression under continuous therapy from under adaptive therapy. The LV model predicts that adaptive therapy delayed the progression of patient #1 by 3.5 months on average (Fig. 3C. continuous (cyan) vs. adaptive (green)) with an average cumulative dose rate of ~46% of continuous therapy. It is worth noting the cumulative dose rate in this study means the percentage of time on therapy, not actual dose, since we are simulating treatment on and off days with the same dose (effect of the drug) per unit time (day). Of note, the one free parameter, the intrinsic growth rate of the sensitive population, was set to be the same as that of the resistant population. The SW model predicts an even more significant improvement from adaptive therapy (~4.6 months) with an average cumulative dose rate of ~53% of continuous therapy (Fig. 3F). We fixed two of the free parameters by setting the growth rates of sensitive cells to be the same as the estimated resistant cell growth rates (*r_s_ = r_R_*), and the transition rate to be 45% of the resistant cell intrinsic growth rate (*β* = 0.45*r_R_*).

### Key parameters that determine clinical gains

Both forms of the mathematical model predicted that adaptive therapy would be beneficial, but the patient-specific benefits were substantially different across the 8 patients (Fig. 4A). The LV model predicted that the average time gained for patient #1 is about 3.5 months, while that for patient #8, is only about one month (Fig. 4A).

**Figure 4.**
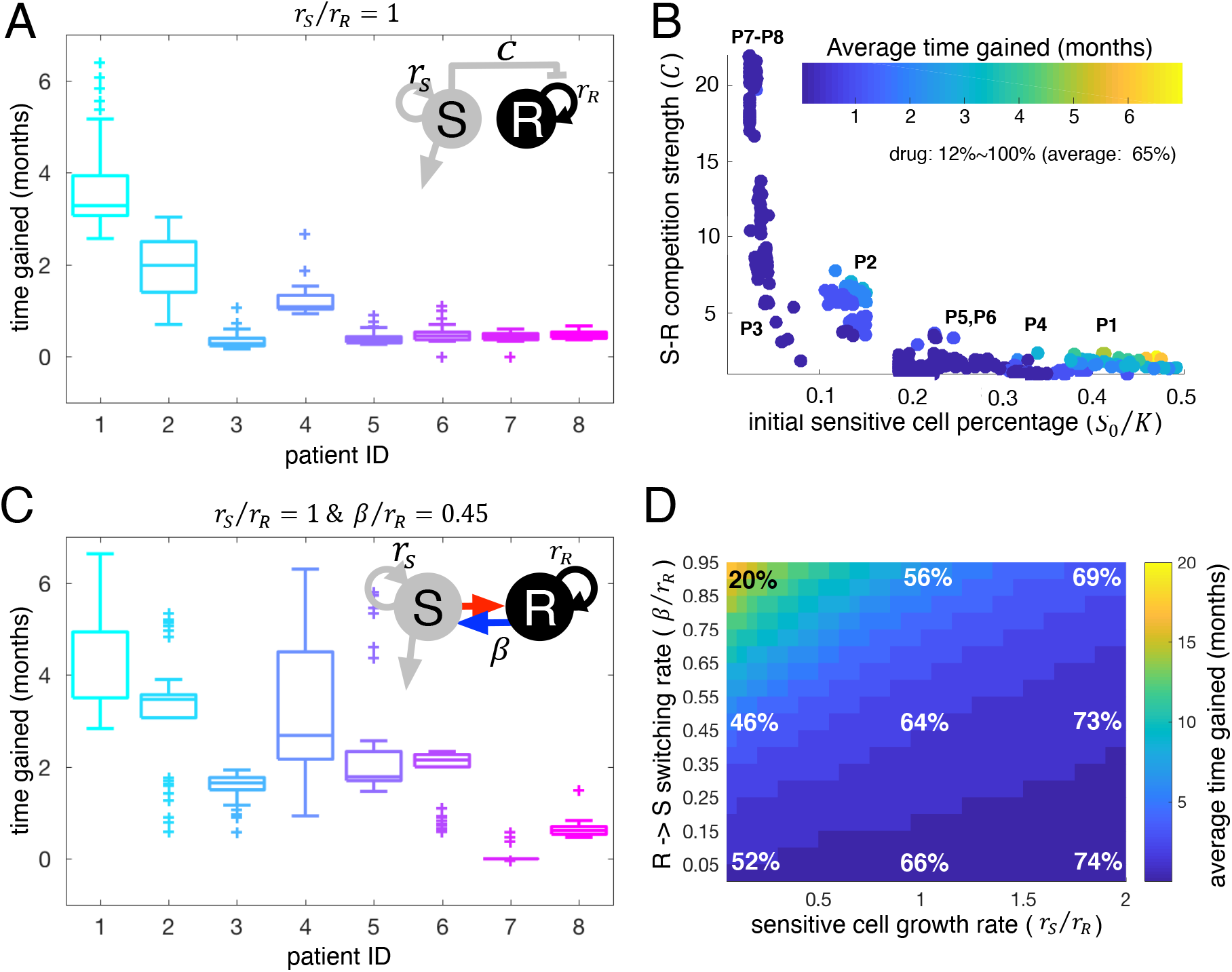
Adaptive therapy simulat i ons. A: The LV model predicted time ga i ned in months with adaptive therapy compared to continuous therapy (time to progression under adaptive therapy - time to progression under continuous therapy) for all 8 patients. All model parameters except the growth rate of sensitive cells are estimated from the LV model fits to individual patient LDH data. The intrinsic growth rate of sensi t i ve cells was set to be the same as the estimated intrinsic growth rate of resistant cells 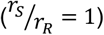. B: Time gained of all 400 virtual patients (50 virtual patients for each actual patient & 8 actual patients) as a function of initial sensitive cell proportion (*S*_0_/*K*) and *S-R* competition coefficient (*C*). Dots: Average virtual patient assuming *r_s_* = 5% – 200% of the estimated *r_R_* per each estimated parameter set obtained from the real patients. C: The SW model predicted time gained in months with adaptive therapy for all 8 patients. All model parameters except for the intrinsic growth rate of sensitive cells (*r_s_*) and *R→S* transition rate (*β*) were estimated from SW model fits to LDH data. The intrinsic growth rate of sensitive cells was set to be the same as the estimated intrinsic growth rate of resistant cells 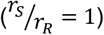 and the *β* was set to 45% of the estimated intrinsic growth rate of resistant cells (*β* = 0.45*r_R_*). D: Average time gained for all 400 virtual patients varied with the intrinsic growth rate of the sensitive cells (*r_s_* = 0.05*r_R_*~2*r_R_*) and the *R→S* transition rate (*β* = 0.05*r_R_*~0.95*r_R_*). Color: time gained in months. Percent represents the average dose rate relative to continuous therapy.

To further explore the significant differences across the 8 patients in the LV model, we simulated patient-specific adaptive therapies using the cohort of 400 virtual patients over a broad range of intrinsic growth rates (*r_S_* = 5% – 200% of the estimated *r_R_*). The estimated competition coefficient, *C*, and the estimated initial size of the sensitive population (*S*_0_/*K*) are key for determining the benefit (time gained) of adaptive therapy relative to continuous therapy (Fig. 4B). In Figure 4B, each colored dot is an average virtual patient of *r_s_* = 5% – 200% of the estimated *r_R_* per each estimated parameter set obtained from the real patients. If the initial sensitive cell percentage is smaller than 10%, time to tumor progression under adaptive therapy is not significantly different from continuous therapy (time gained ≤ 1 month, patient #3, #7, #8). The larger the competition coefficient (C), the more time gained (Fig. 4B), provided that the initial sensitive cell population ranges from 10% to 40% of carrying capacity. Adaptive therapy is most successful for tumors with a high initial number of sensitive cells (> 40% of *K*, see patients #1). For adaptive therapy, the percent time on treatment varied between patients (i.e., the cumulative dose rate of 12% to 100% of continuous therapy), with an overall average of 65%. In the simulations, we assume treatment dose per treatment day in adaptive therapy is the same as continuous therapy. The drug dose implies time on the drug. The cumulative dose rate of 12% means 12% of the time on the drug compared to continuous therapy.

To explore the significant differences of adaptive therapy response across the 8 patients in the SW model, we simulated adaptive therapy using the cohort of 400 virtual patients generated from fitting the SW model to the patient data (Fig. 1 & Fig 3D). Adaptive therapy is predicted to delay progression even more (Fig. 4C vs. Fig. 4A) with this model. Among the 8 patients, adaptive therapy increased patient #1 ‘s time to progression by up to 7 months while patient #8 still gained only one month on average when we fixed two of the free parameters *r_s_*(= *r_R_*) and *β*(=.45*r_R_*). We further ran a suite of simulations with the virtual cohort and compared the average time gained. It is worth noting again that most model parameters were estimated from each patient LDH (Fig. 1A) except for two parameters, the intrinsic growth rate of sensitive cells (*r_s_*) and the switching rate from resistant to sensitive during treatment holidays (*β*). For these two free parameters (*r_s_, β*), we considered a range of values (*r_s_* ∈ [.05*r_R_*, 2*r_R_*], *β* ∈ [.05*r_R_*, .95*r_R_*]). A larger switching rate (*β*) results in a larger time gained from adaptive therapy (Fig. 4D). This effect occurred for all patients. Interestingly, changing the intrinsic growth rate of sensitive cells had little impact on the efficacy of adaptive therapy if the switching rate is low (time gained ≤ 2 months for all *r_s_* if *β* < .1*r_R_*). If the switching rate is larger than 0.1*r_R_*, a tumor composed of slower growing sensitive cells responded better to adaptive therapy (increasing time gained relative to continuous therapy). The cumulative dose rates of adaptive therapy varied substantially over the ranges of parameters. For large switching and slow-growing tumors, the cumulative dose rate was 20% of continuous therapy, while for low switching rates and fast-growing tumors, the rate was 74% of continuous therapy.

Both model formulations demonstrate that, in general, adaptive therapy can be effective in delaying tumor progression using significantly less drug than continuous therapy. For the LV model, adaptive therapy is most effective if tumors are composed of a sufficiently large number of sensitive cells initially (>40% of carrying capacity *K*). Adaptive therapy gains in superiority as the sensitive cells have a higher competition coefficient (*C*) and exert more significant inhibition on resistant cells through competition. The SW model highlights that adaptive therapy could be beneficial if drug-sensitive and -resistant states are phenotypically plastic. The predicted time gained is about 20 months if the switching rate from drug-resistant to drug-sensitive cells is large, and the sensitive cell growth rate is slow during treatment holidays. Even if the switching rate is very low, adaptive therapy can delay time to progression from 2-5 months. Taken together, our simulations show the potential benefits of adaptive therapy and identify the key parameters and conditions for adaptive therapy to be superior in controlling tumor volume relative to continuous therapy.

### A different treatment-stopping criterion

So far, for adaptive therapy, we used 50% of the initial tumor burden as the treatment-stopping criterion. This criterion was implemented in the first clinical trial of adaptive therapy for castrate-resistant prostate cancer (20). Would a less aggressive treatment stop criterion be better as suggested by more recent models (27, 28, 34)? We simulated a suite of adaptive therapy simulations with a new stopping criterion of 20% reduction to address this question. Therapy is held off if the tumor burden decreases to below 20% of the initial burden. The tumor is rechallenged with therapy when the burden returns to its initial value.

The LV model predicts that a 20% stopping criterion is indeed a better threshold if tumors are already responding well to adaptive therapy. In Figure 4B, we showed that tumors respond well if the initial population size of sensitive cells is high, and the competitive effect of sensitive on resistant cells is large. Under these conditions, the 20% threshold is more effective in controlling tumor burden than the 50% threshold (Fig. 5A, Patient #1 under the 20% threshold adaptive therapy gained 7-20 months over continuous therapy vs. Fig. 4B, Patient #1 under the 50% threshold gained 3-7 months over continuous therapy). We observed a similar pattern in the SW model. A lower threshold is more effective if tumors are already responding well. If the switching rate from resistant to sensitive cells is large and the intrinsic growth rate of sensitive cells is small, adaptive therapy with either the 50% or 20% threshold substantially delays time to progression relative to continuous therapy. Under these conditions, an adaptive therapy with the 20% threshold delays progression up to 5 months more (up to 25 months gained Fig. 5B vs. up to 20 months gained with 50% threshold Fig 4D). Taken together, a smaller tumor burden reduction criterion may be more effective in delaying tumor progression when the tumors underlying dynamics satisfy specific conditions, such as a large number of initially sensitive cells, strong competitive inhibition of resistant cells by sensitive ones, a high switching rate from *R* to *S*, and a slow-growing sensitive population.

**Figure 5.**
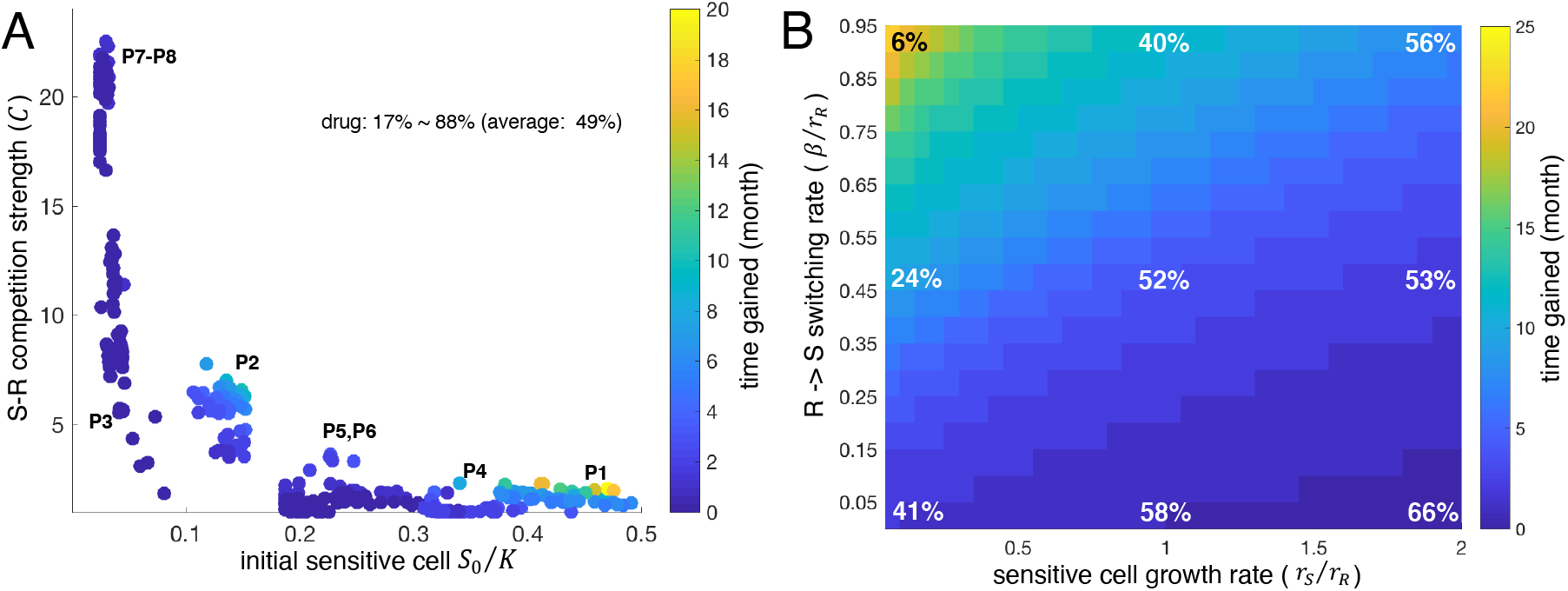
Adaptive therapy simulation with a 20% burden decrease as a stop-treatment criterion. A: Average time gained in months with 20% adaptive therapy compared to continuous therapy (time to progression on 20% threshold adaptive therapy – time to progression on continuous therapy) predicted by the LV model as a function of initial sensitive cell proportion and *S-R* competition coefficient. B: Average time gained predicted by the SW model as a function of sensitive cell growth rate and *R→S* transition rate. Percent represents the average dose rate relative to continuous therapy.

### Progression free survival

Finally, we compared the probability of progression free survival of all virtual patients under continuous, −50% and −20% stopping adaptive therapies. Since we developed two mathematical models, LV and SW, we generated two sets of Kaplan-Meier curves (Fig. 6, A: LV, B: SW). For these K-M curves, we utilized all virtual patients with either the LV or the SW model. This “trial” group was then subjected to the 2 adaptive trial procedures (−50% and −20% stopping threshold) and continuous therapy. Here, we assume that tumor progression occurred when LDH levels reached 150% of their initial LDH (Fig. 6, A: LV, B: SW). The two models predicted a longer progression free survival under both adaptive therapies (p < 0.001), with a 20% threshold being superior to the 50% stopping criterion.

**Figure 6:**
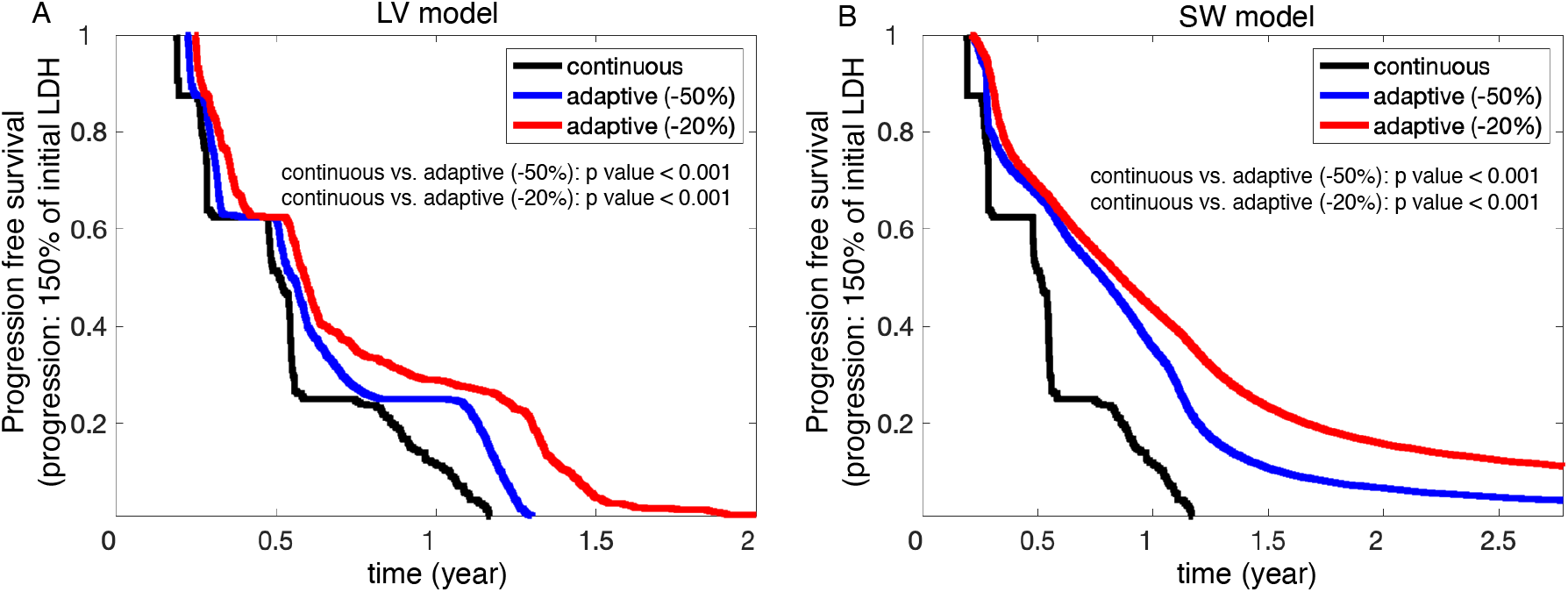
Progression free survival comparison. A. The LV model predicted progression free survival under continuous therapy (black) and two different adaptive therapy stopping thresholds (−50%: blue & −20%: red). The survival probabilities are significantly different (p-values < 0.001). B. The SW model predicted progression free survival under continuous (black), −50% adaptive (blue), and −20% adaptive (red) therapies. The survival probabilities are significantly different (p-values < 0.001)

## Discussion

The adaptive abiraterone trial for metastatic castration-resistant prostate cancer (NCT02415621) extended time to progression over 27 months (20). The success of this trial has inspired three more clinical trials testing for the feasibility of adaptive therapy in patients with castrate sensitive prostate cancer (NCT03511196), advanced melanoma (NCT03543969), and thyroid cancer (NCT03630120). The actual real-world benefit from adaptive therapy would likely vary among patients, as already observed in (20). Understanding the underlying mechanism for this variability may be crucial for patient selection in future clinical trials. As previous mathematical models have demonstrated (16, 17, 20, 23, 24), the interaction between the drug-sensitive population and resistant population drives the outcome of adaptive therapy. These competitive aspects have been further investigated experimentally (35), but more work needs to be done to better understand how resistance emerges and is maintained under different treatment strategies. A sensitive cell population can inhibit the growth of a resistant population via competition. Sensitive cells can acquire resistance, and resistant cells can be re-sensitized to the drug. To examine these interactions in the context of metastatic melanoma, we developed two different mathematical models, a standard Lotka-Volterra competition model (LV) with a varying competition coefficient (effect of sensitive cells on resistant) and an extension with phenotype switching (SW) but with all competition coefficients set equal to 1. The LV model describes the competition between distinct sensitive and resistant populations for a limited resource. In contrast, the SW model assumes individual cells can acquire resistance when therapy is on and become re-sensitized when therapy is off.

Our analyses and simulations of these two models demonstrate that adaptive therapy, with timed treatment holidays, prolongs survival compared to standard of care (MTD continuous therapy). We identified key tumor parameters where this benefit is most significant. The LV model shows that adaptive therapy can extend the time to progression significantly whenever the initial population of sensitive cells is large enough, and when sensitive cells have a large competitive effect on resistant cells. The SW model illustrates that adaptive therapy is most beneficial whenever the re-sensitization rate of the resistant cell population to the sensitive population is high, and the growth rate of sensitive cells is low. Under these constraints, adaptive therapy can more than double the time to progression (max time of delay is 20 months in adaptive vs. 6 months with continuous therapy). Interestingly, tumors without these properties did not respond well to the standard of care either (< 3 months of progression free survival). In these cases, switching to alternative therapies as soon as possible might be more desirable. For example, in metastatic melanoma, adding or switching to immunotherapy may provide a clinical benefit, particularly for patients expected to respond poorly to either adaptive therapy or continuous therapy. Taken together, our study has identified a patient group who may benefit the most from adaptive therapy, with predicted clinical time gains for such patients. This study raises the challenge and the opportunity of directing clinical research towards swiftly measuring the key parameters for a given patient to facilitate truly personalized medicine and deciding on the efficacy of adaptive therapy.

A key assumption of our analysis is that a patient initial tumor burden is tolerable and not immediately life threatening. Maintaining a potentially large tumor burden, as opposed to focusing on shrinking tumors as fast as possible, can make both clinicians and patients uneasy. There are cancers where an initial burden is not tolerable and may be life-threatening (e.g., aggressive glioblastoma or very painful tumors that may be bleeding, etc.). There are, however, situations where a patient initial tumor burden is tolerable, and the patient is asymptomatic. In fact, maintaining the sum of all tumor diameters (tumor burden) is one of the response evaluation criteria in the measurement of solid tumors in clinical trials. Stable disease is defined as a < 20% increase in the sum of target lesions per the widely used RECIST 1.1 criterion (Response Evaluation Criteria in Solid Tumors) (36). There is increasing evidence that chronic control of disease burden is more effective in improving patient survival, not only in cancers (20) but also in bacterial infections (37-40). Our proof of concept analysis on patients with advanced melanoma treated with BRAF/MEK inhibitor targeted therapy illustrates that maintaining a tolerable tumor burden may delay progression significantly.

The mathematical models presented here are simplified representations of what may be happening in actual cancers under treatment. The models rest on the assumption that two key cancer cell populations compete and interact with each other in a well-mixed environment. In reality, a tumor is not a well-mixed population of cancer cells but is a spatially stratified population (41-44) that drives different pharmacokinetics (45) and is impacted by the influence of a heterogeneous and dynamic microenvironment (46-49). Such aspects could be incorporated in future studies, however, adding more complexity does not guarantee a better understanding. For now, the clinical data is non-spatial (blood levels of LDH). Therefore, while a spatial model may deliver a better representation of the tumor in a more relevant context, it would be more complex and burdened with additional assumptions that cannot be evaluated by blood biomarkers alone.

We chose our modeling approach as a starting point to better understand how different modes of drug-sensitive and -resistant interactions, distinct sensitive and resistant subpopulations vs. phenotypic switching, impact the outcomes of adaptive therapy. The results presented show the need to better understand the properties of the sensitive and resistant populations. If they are genetically distinct and preexisting, then for our cohort of 8 patients, adaptive therapy would provide less benefits to fewer of the patients than if drug sensitivity and resistance are reversible phenotypically plastic states. These properties of the sensitive and resistant populations must be determined empirically. We have previously examined the behavior of the metastatic melanoma cell line WM164 and found it spears to exhibit this more phenotypically plastic behavior, where as another melanoma cell line 1205LU did not (23). Therefore, there may be significant heterogeneity across patients in terms of which resistance mechanisms are at play and how the emerge and are maintained.

Equally as important as the phenotypic heterogeneity is the actual tumor burden - accurate and frequent measurement of tumor burden is key to calibrating mathematical models so that they can better reproduce tumor dynamics and make reliable predictions. In the ongoing adaptive therapy prostate cancer trials, prostate-specific antigen (PSA) is used as a marker for disease burden (NCT02415621 & NCT03511196). LDH is a standardized systemic biomarker for melanoma, although it can also increase in certain conditions not related to tumor burden, such as in patients who have liver toxicity from drug treatment (50). Newer blood biomarkers such as circulating tumor DNA levels may correlate better with tumor response and progression in melanoma patients (51). These biomarkers, however, may only highlight the sensitive cell population and all of them obviously lack spatial information. Development of novel imaging technologies is urgently needed to allow for non-invasive serial assessment of tumor burden; not only would this provide greater temporal resolution but would offer the opportunity to better understand the spatial dynamics.

Our analysis of patients with advanced melanoma identifies tumor-specific conditions (parameters) and resulting dynamics where adaptive therapy may lead to significant clinical gains. Identifying such patients before treatment would accelerate clinical translation. Here, we identify such parameters by calibrating our models to individual patient tumor burden dynamics. This study highlights another potential benefit of using mathematical models in the clinic by supporting patient selection in clinical trials based on tumor parameter identification. Having chosen the appropriate patients, the mathematical models can then be tailored to each individual patient treatment response to predict and drive their next treatment decision. This mathematical model driven treatment decision paradigm is especially critical for adaptive therapy (and more generally, personalized medicine), since it directly facilitates both therapy dosing and switch points. We, therefore, advocate for a greater integration of predictive mathematical models in the clinical decision process.

## Supplementary material

**Figure S1.**
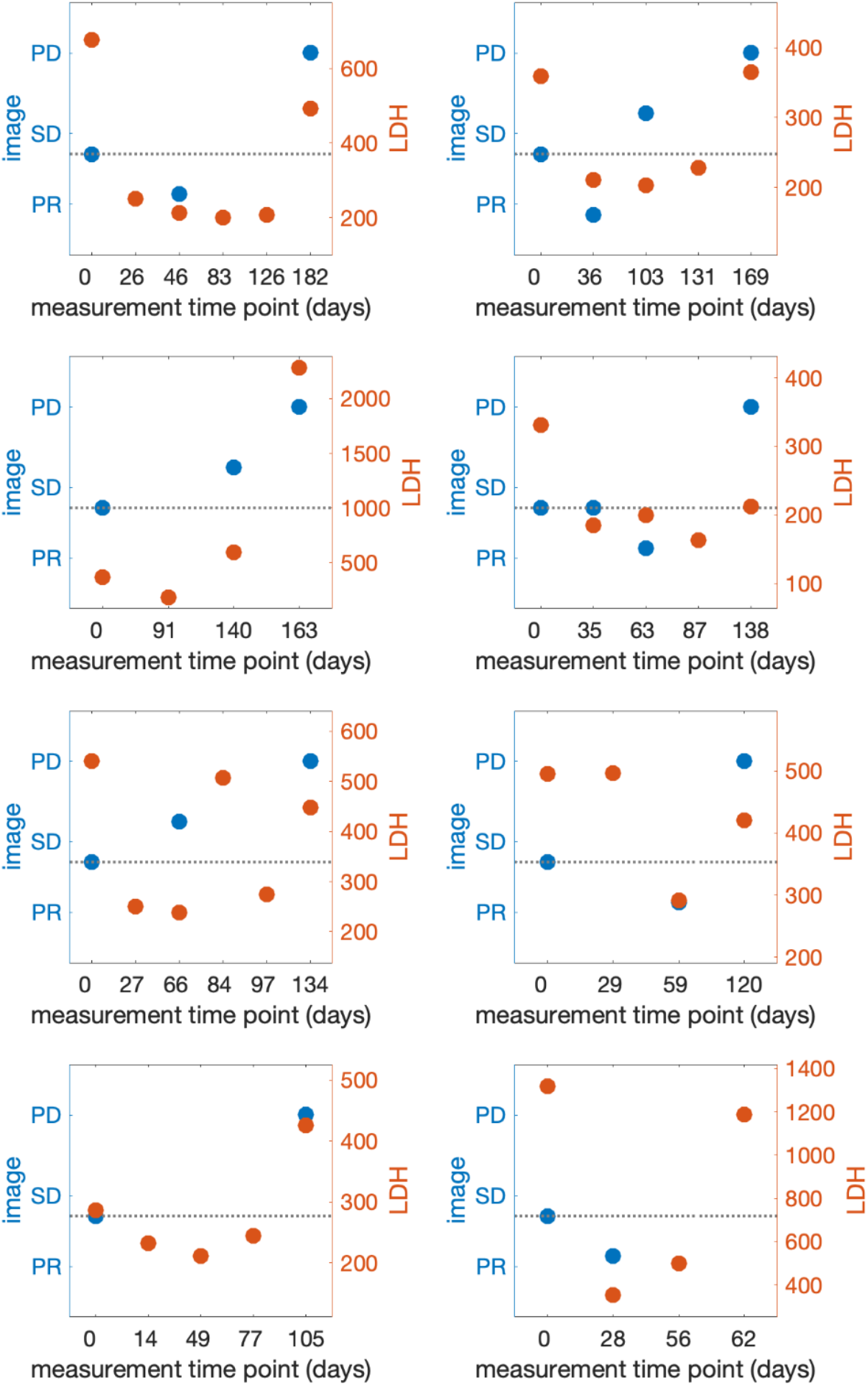
Patient LDH and image (CT) data. Partial Response (PR): more than 30% reduction in the sum of target tumor diameters, Stable Disease (SD): up to 20% increase in the sum of target tumor diameters, Progressive Disease (PD): more than 20% increase in the sum of target tumor diameters. Blue: CT image (left axis), Orange: LDH (right y-axis).

**Table S1.**
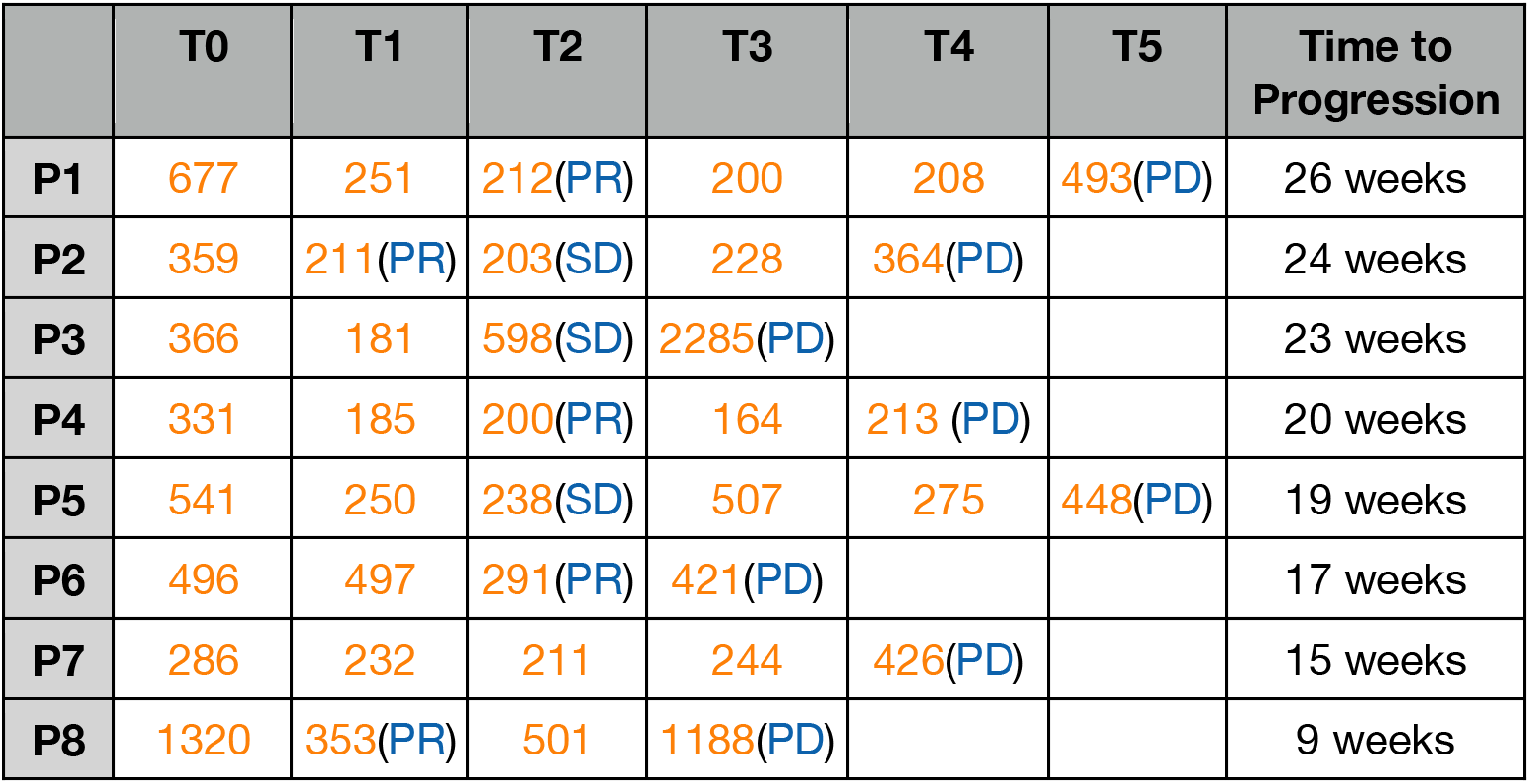
Patient LDH (yellow) and response evaluation data (blue). T0-T5: measurement time.

**Figure S1.**
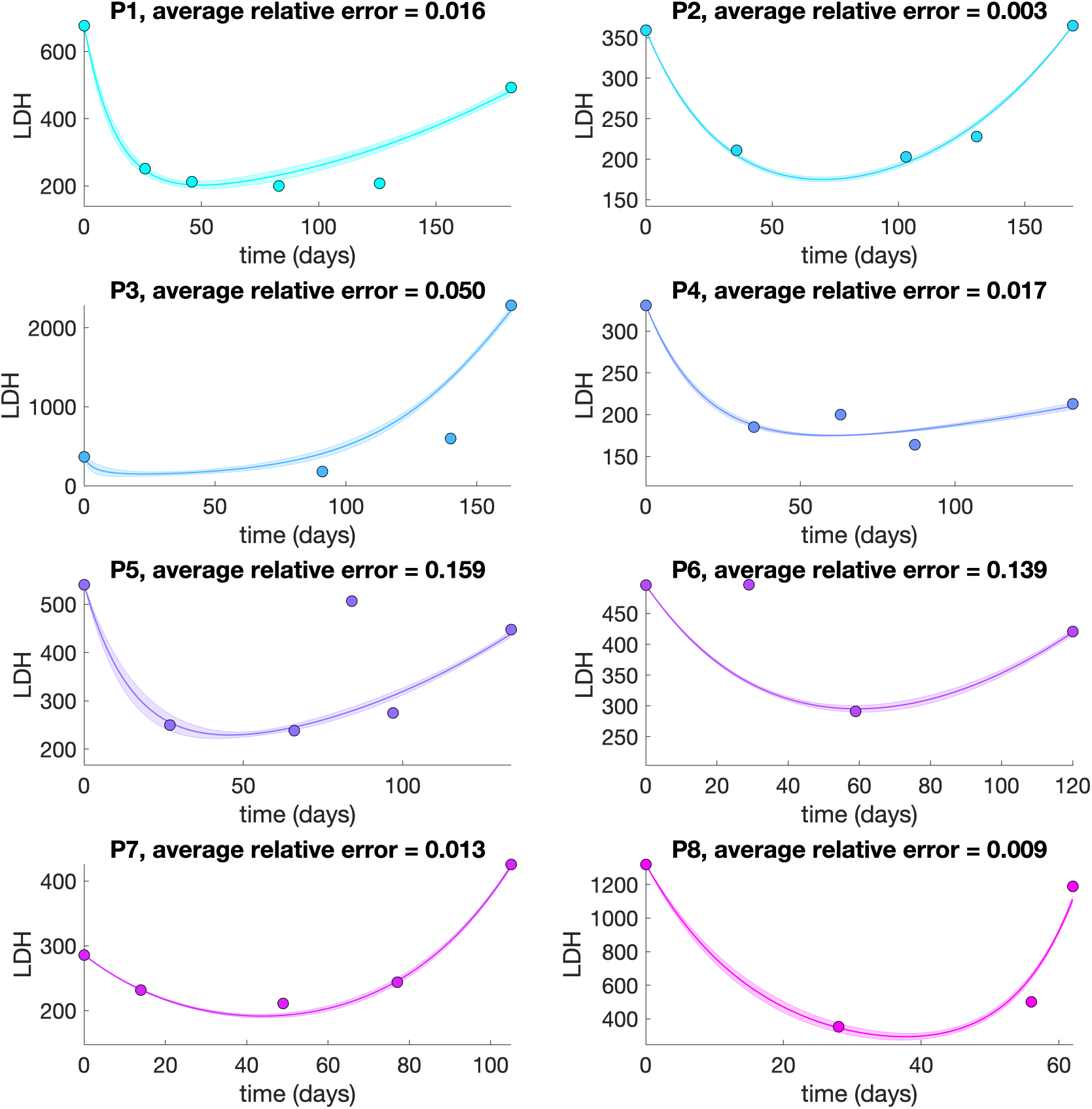
LV model calibration. We estimated cell growth rates, carrying capacity, drug-sensitive population death rate, and competition coefficient. The model parameters were estimated by a method described in the Parameter Estimation section. In brief, we estimated model parameters that minimized the difference between model predicted LDH and each patient LDH. The implicit filtering algorithm with different initial values was used to identify various sets of parameters that equally fit to patient data. We selected the top 50 best-matched parameters to generate fitted curves. Thick line: mean of model predictions, shadow: standard deviation, dot: patient LDH measurement.

**Figure S2.**
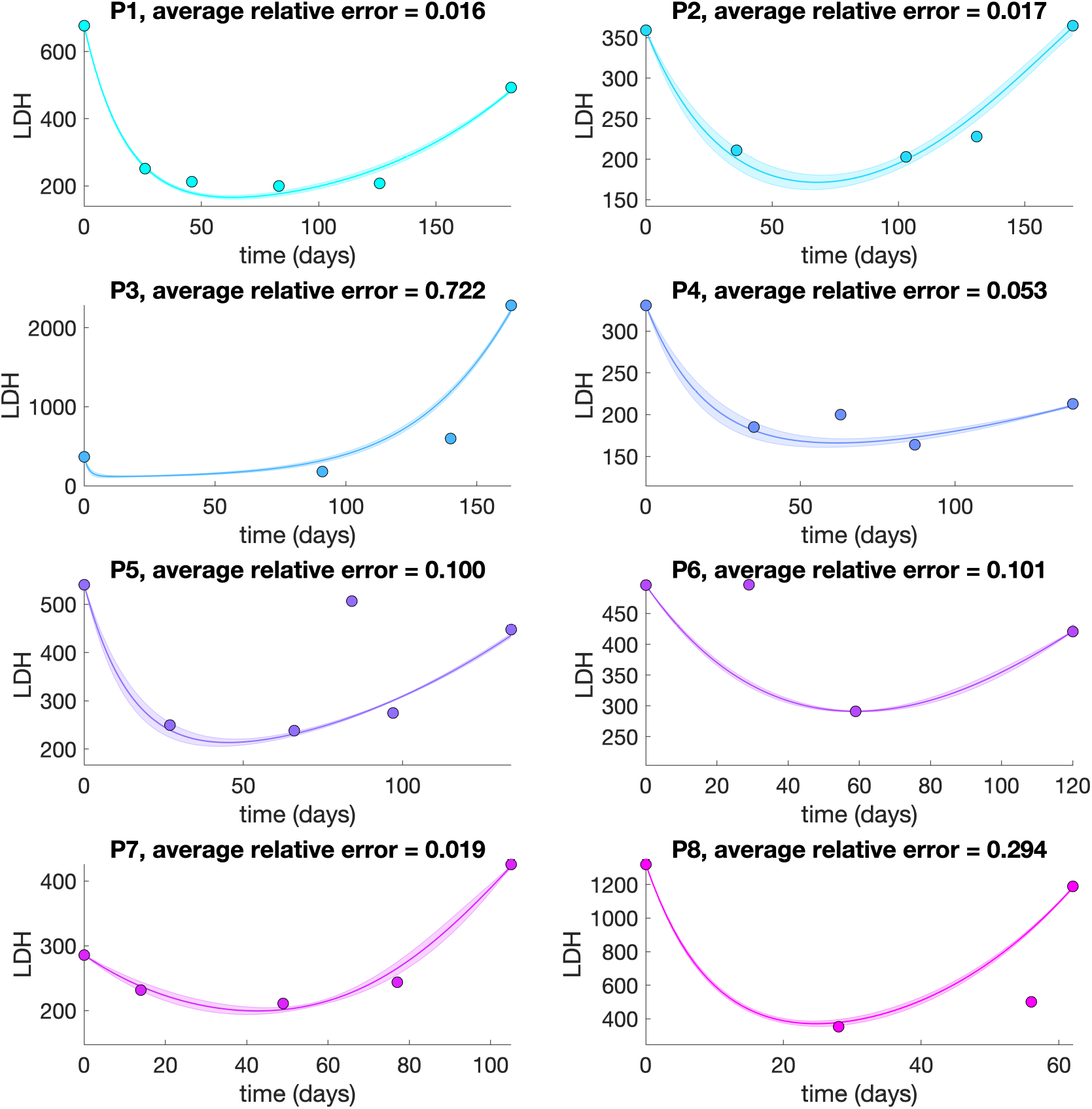
SW model calibration. For SW model, we estimated growth rates, carrying capacity, and transition rates. Thick line: mean value of model predictions, shadow: standard deviation, dot: patient LDH data.

## Notes

### Competing Interest Statement

Eunjung Kim, Joel S. Brown, Alexander R.A. Anderson: no conflict of interest to declare. Zeynep Eroglu: advisory boards with Array, Genentech, Novartis, Regeneron, Natera, SunPharma, and research funding from Novartis.

